# Generation of conditional auxin-inducible degron (AID) cells and tight control of degron-fused proteins using the degradation inhibitor auxinole

**DOI:** 10.1101/530410

**Authors:** Aisha Yesbolatova, Toyoaki Natsume, Ken-ichiro Hayashi, Masato T. Kanemaki

## Abstract

Controlling protein expression using a degron is advantageous because the protein of interest can be rapidly depleted in a reversible manner. We pioneered the development of the auxin-inducible degron (AID) technology by transplanting a plant-specific degradation pathway to non-plant cells. In human cells expressing an E3 ligase component, OsTIR1, it is possible to degrade a degron-fused protein with a half-life of 15–45 min in the presence of the phytohormone auxin. We reported previously the generation of human HCT116 mutants in which the C terminus of endogenous proteins was fused with the degron by CRISPR–Cas9-based knock-in. Here, we show new plasmids for N-terminal tagging and describe a detailed protocol for the generation of AID mutants of human HCT116 and DLD1 cells. Moreover, we report the use of an OsTIR1 inhibitor, auxinole, to suppress leaky degradation of degron-fused proteins. The addition of auxinole is also useful for rapid re-expression after depletion of degron-fused proteins. These improvements enhance the utility of AID technology for studying protein function in living human cells.

## 1. Introduction

Conditional depletion of a protein of interest (POI) is a powerful approach to analyse its function *in vivo*, especially for POIs that are essential for cell viability. Recently, conditional approaches using a degron have been drawing increased attention [1]. A degron-fused protein can be rapidly and efficiently degraded when needed, so that the primary defect arising from the depletion can be observed before the phenotype is complicated or compromised by secondary defects. For this purpose, we pioneered the establishment of the auxin-inducible degron (AID) technology to control degron-fused proteins in yeast and mammalian cells (**Figure 1A**) [2]. When expressed in non-plant cells, TIR1 of rice (OsTIR1) forms a complex with the endogenous SCF (Skp1–Cul1–F box) components. The SCF–OsTIR1 E3 ubiquitin ligase is only activated when IAA or NAA (a natural or synthetic auxin, respectively) is bound (**Figure 1A**). We identified a 7 kD degron termed mini-AID (mAID) and others identified similar AID degrons (**Supplementary Figure 1**) [3–5]. A POI fused with an mAID is recognized by SCF–OsTIR1 for ubiquitylation and subsequent proteasomal degradation (**Figure 1A**).

**Figure 1.**
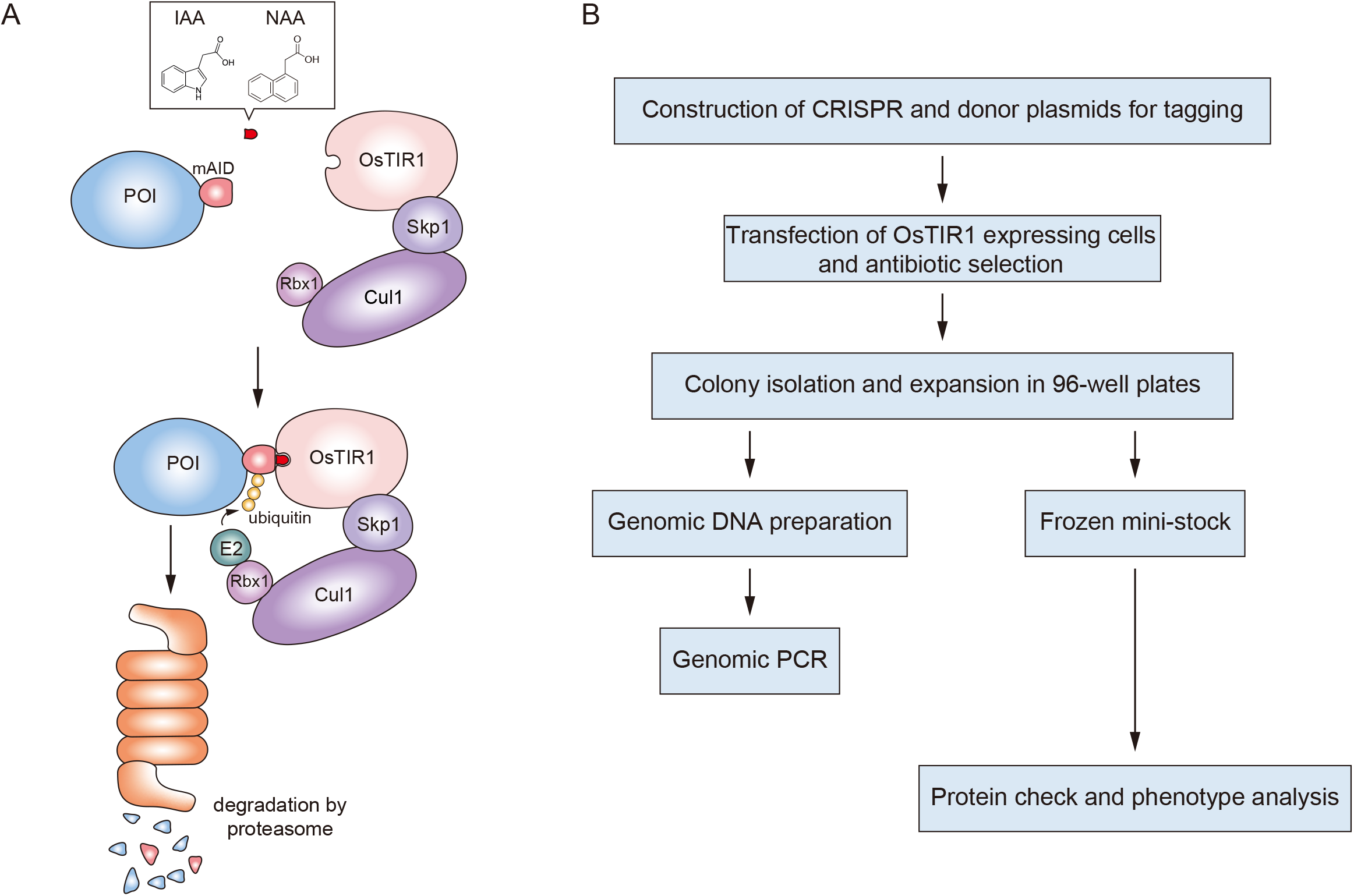
Schematic illustration of the AID system. (**A**) Indole-3-acetic acid (IAA)- or 1-naphthaleneacetic acid (NAA)-bound OsTIR1 associates with mini-AID (mAID) and promote ubiquitylation of the mAID-fused protein for proteasomal degradation. (**B**) The strategy used for generating conditional AID cells by tagging an endogenous POI.

Recently, we showed that conditional human cells can be generated by tagging endogenous genes with an mAID cassette using CRISPR–Cas9-based gene tagging [6]. This technology has recently been applied in many studies. One particular example is the assessment of chromosome architectures [7–10]. Moreover, the AID technology has been applied to other model organisms, such as fission yeast, fruit fly, nematode, zebrafish and the parasitic *Toxoplasma gondii* [11–15]. These studies support the idea that the AID technology can be used as a standard method to achieve conditional protein depletion. However, a drawback of this technology is the reduced expression level of mAID-fused target proteins in OsTIR1-expressing cells [6]. This “basal degradation” might be caused by the presence of contaminating auxin-like chemicals in bovine serum or culture media [1]. To achieve tight control of the expression of mAID-fused proteins, an improvement in this technology would be useful.

Here, we describe a method to generate conditional human HCT116 and DLD1 mutants by homology-directed repair (HDR)-mediated gene tagging using CRISPR–Cas9 (**Figure 1B**). We offer a new series of plasmids for N- or C-terminal tagging with mAID and other tags. To overcome the problems associated with basal degradation, we used a TIR1 inhibitor called auxinole [16]. It was possible to suppress basal degradation and rapidly recover expression after depletion by supplementing culture media with auxinole.

## 2. Construction of CRISPR and donor plasmids for tagging

Figure 1B shows the procedures that were used to generate conditional AID cells, which typically require one month of work. We previously reported mAID tagging at the C terminus of a POI (**Figure 2A**) [6]. We now developed a procedure to tag a POI with mAID at the N terminus (**Figure 2B**) [17]. We offer C- or N-terminal tagging plasmids with mAID or other tags at Addgene (https://www.addgene.org/Masato_Kanemaki/) and the National Bio-resource Center (NBRP) (http://dna.brc.riken.jp/en/gsb0000en/rdb08468) (**Figure 2C**).

**Figure 2.**
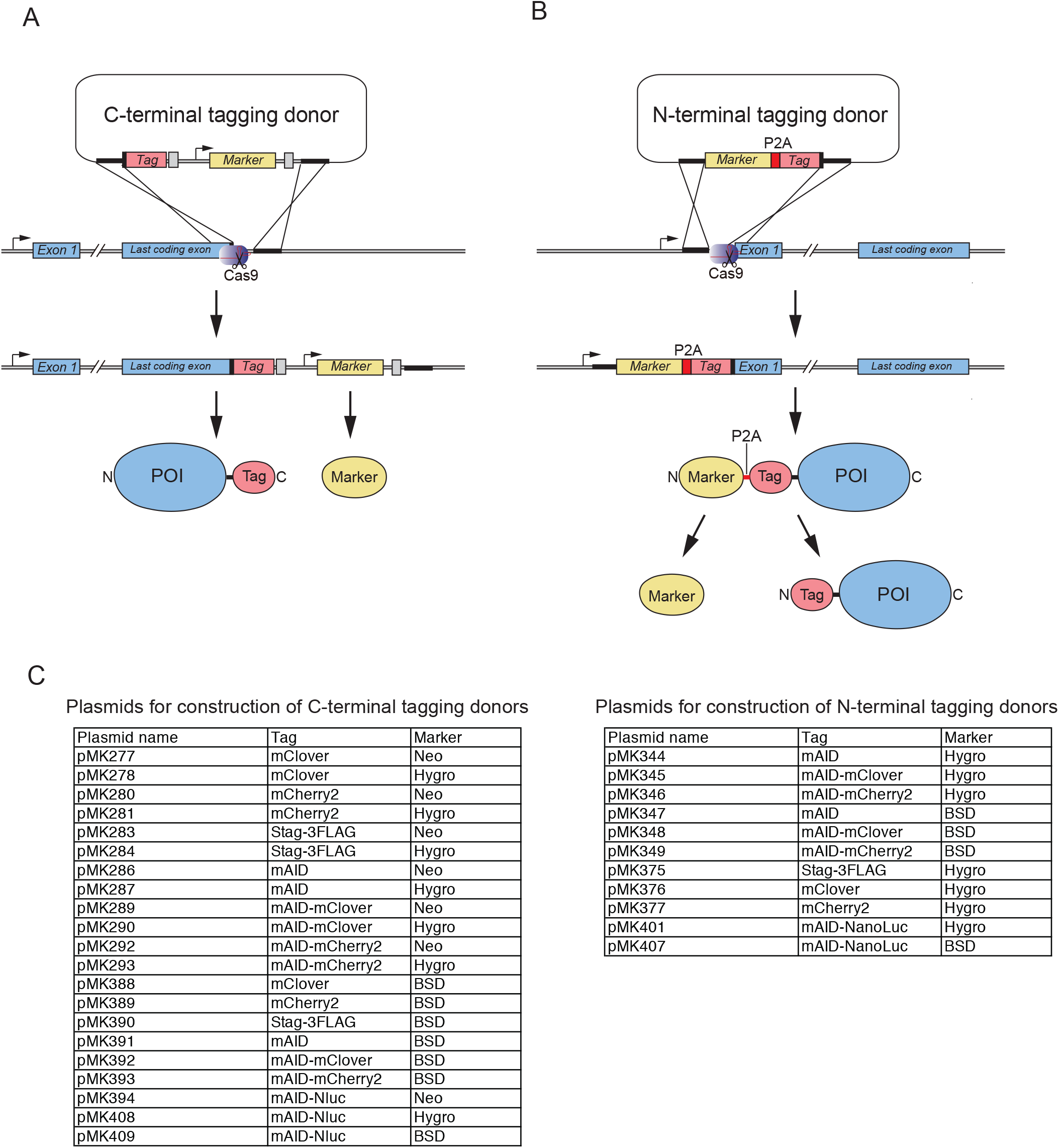
Tagging of an endogenous POI using CRISPR–Cas9-based genome editing. (**A**) Fusing a tag at the C terminus of a POI. CRISPR–Cas9 generates a double-strand break (DSB) near the stop site for insertion of a donor harbouring a tag and a marker. The fusion protein and the marker will be expressed independently. (**B**) Fusing a tag at the N terminus of a POI. CRISPR–Cas9 generates a DSB near the first ATG site for insertion of a donor harbouring a marker-P2A-tag cassette. The fusion proteins will be processed at P2A to express the marker and the tag-fused POI. (**C**) List of N- and C-terminal tagging plasmids.

### 2.1. Construction of a CRISPR–Cas9 plasmid

To identify a CRISPR–Cas9 targeting site, we usually choose an appropriate sequence within 50 bp upstream or downstream from the ATG or stop codon. We use the following target finder sites.

- IDT custom Alt-R guide design: https://sg.idtdna.com/site/order/designtool/index/CRISPR_CUSTOM
- WEG CRISPR finder: https://www.sanger.ac.uk/htgt/wge/

We mainly use pX330-U6-Chimeric_BB-CBh-hSpCas9 (Addgene #42230) to express the SpCas9 nuclease and guide RNA according to the protocol of Ran et al. [18]. However, it should be possible to use a plasmid encoding other Cas9 variants. As discussed in the next section, it is important to destroy or lose the target site within the genome by the HDR-mediated insertion of a tagging cassette.

### 2.2. Construction of a donor plasmid

We described previously a method to construct donor plasmids by generating homology arms (HAs) using long primers and gene synthesis [6]. A downside of this strategy is that the HAs are relatively short (up to 200 bp each) and that it can be costly. As an alternative (and economic) approach, we describe a method to clone HAs from the genomic DNA (**Figure 3**). The C- or N-terminal coding region (about 1000 bp) is amplified by PCR, followed by cloning into a conventional cloning plasmid (such as pBluescript II). After confirming the sequence, a cloning site for inserting a tagging cassette with a selection marker is created by inverse PCR. The cassette is cloned at the cloning site to complete the construction of the donor plasmid, which contains the HA at both ends (about 500 bp each) (**Figure 3A and B**). Importantly, the donor plasmid has to be designed to destroy the CRISPR target site when inserted into the genome. In the case of targeting a noncoding locus, it is possible to delete part of the target sequence, unless a functional noncoding element is absent. In the case of targeting a coding locus within a gene, it is important to introduce silent mutations, to avoid re-cutting after HDR-mediated insertion. Another important point is that a cassette encoding a tag has to be cloned in frame with the gene of interest, to be able to express a fusion protein. All cloning procedures can be carried out using a standard cloning method. In cases in which standard cloning using restriction enzymes and ligase is difficult, it is possible to employ a recombinase-mediated cloning method, such as In-Fusion cloning or Gibson assembly.

**Figure 3.**
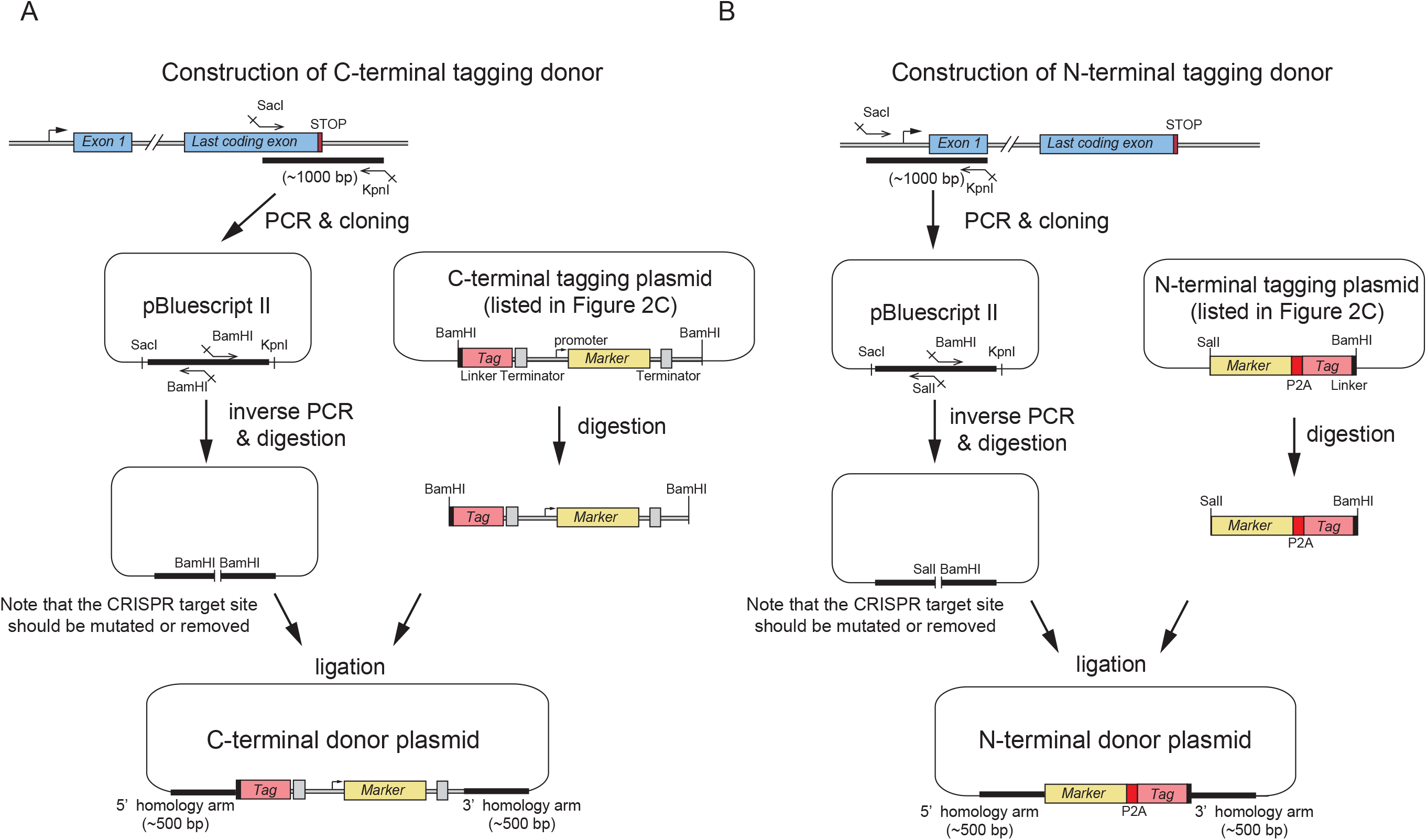
Schematic illustration of donor plasmid construction. (**A**) A DNA fragment containing the stop site (about 1 kb) is cloned into a cloning plasmid (such as pBluescript II). After creating a restriction enzyme site (such as BamHI) by inverse PCR, a DNA fragment containing a tag and a marker is cloned into the plasmid, to generate a donor vector. (**B**) A DNA fragment containing the first ATG site (about 1 kb) is cloned into a cloning plasmid (such as pBluescript II). After creating restriction enzyme sites (such as SalI and BamHI) by inverse PCR, a DNA fragment containing marker-P2A-tag is cloned into the plasmid, to generate a donor vector.

To select clones with a biallelic insertion efficiently, we transfected two donor plasmids containing a Neo or Hygro marker, respectively, for dual-antibiotic selection [6]. As an alternative approach, it is possible to generate clones with a biallelic insertion using a single donor plasmid with a single selection marker. In this case, both monoallelic and biallelic clones can be obtained.

## 3. Transfection and colony formation

We established near-diploid colon cancer cell lines, HCT116 and DLD1, constitutively or conditionally expressing OsTIR1 (CMV- or Tet-OsTIR1, respectively) by inserting the transgene at the safe AAVS1 locus [6]. These parental cell lines are available from NBRP or our laboratory upon request. Alternatively, it is possible to generate a new parental cell line. In this case, the plasmids that can be used for introducing OsTIR1 at AAVS1 are available at Addgene and NBRP (**Supplementary Figure 2**) [6]. A CRISPR–Cas9 plasmid for targeting the AAVS1 locus is also available from Addgene (AAVS1 T2 CRISPR in pX330: #72833).

### 3.1. Transfection

1. Prepare a sub-confluent culture of low-passage-number HCT116 or DLD1 CMV/Tet-OsTIR1 cells.
2. Collect cells by trypsinization to prepare a cell suspension. Count and dilute the cells to 1 × 10^5^ cells/mL. Seed 1 mL of the diluted cells per well in a 12-well plate and culture the cells at 37 °C for two days. HCT116 cells are slow to attach to the bottom. We allow two days of culture before transfection.
3. Prepare the transfection mixture by mixing 2 μL of 200 ng/μL CRISPR–Cas9 plasmid or TE (control), 2.5 μL of 200 ng/μL donor plasmid, 45.5 μL of Opti-MEM I Reduced Serum Medium (ThermoFisher Scientific, 31985062) and 4 μL of FuGENE HD Transfection Reagent (Promega, E2311), and incubate the mixture at RT for 15 min before applying to the cells.
4. One day after transfection, collect the cells in 1 mL of the medium after trypsinization. Depending on the efficiency of CRISPR–Cas9 and donor insertion, the number of colonies obtained will vary significantly. To pick up single colonies successfully, prepare several dilutions of transfected cells in a 10 cm dish (dilutions range from 100 to 1000 times; 100 to 10 μl of cell suspension into one 10 cm dish containing 10 mL of culture medium).

### 3.2. Antibiotic selection and colony formation

The final concentration of antibiotics is shown below.

- Neo: 700 μg/mL of G418 Sulfate (potency-based)
- Hygro: 100 μg/mL of Hygromycin B Gold (InvivoGen, #ant-hg)
- BSD: 10 μg/mL of Blasticidin S Hydrochloride

If using a donor harbouring Neo, add G418 soon after plating the transfected cells. Hygromycin B Gold or Blasticidin S should be added one day after plating for cell recovery. In case of dual selection, add G418 and Hygromycin B Gold at the same time one day after plating. Replace culture media with a fresh one containing an appropriate antibiotic every 3–4 days until colonies become visible (usually 11–13 days after plating).

## 4. Colony isolation, expansion in 96-well plates and genotyping

Visible colonies should form on the selection dishes after 11–13 days of culture. If the tagging worked well, you should find more colonies on the plates containing cells transfected with the CRISPR–Cas9 plasmid than in those containing cells without the CRISPR–Cas9 plasmid. Although this is a good indicator of successful tagging in many cases, gene tagging sometimes works well even if no differences in colony number are observed. We usually pick at least 32 and 16 clones for single- and dual-antibiotic selection, respectively, to obtain clones harbouring the tag on both alleles.

### 4.1. Colony isolation

1. Prepare a 96-well plate containing 10 μL of trypsin/EDTA solution in each well.
2. Replace the culture medium in the dish with PBS.
3. Under a stereo microscope, pick a single colony using a micropipette with the volume set at 25 μL (Use a micropipette such as a Gilson pipetman P200.).
4. Transfer the colony to the 96-well plate containing trypsin/EDTA for trypsinization. After transferring each set of 16 colonies, add 100 μL of media to each well, to quench trypsin. Repeat this step to obtain a sufficient number of clones.
5. After isolating clones, resuspend cells in the culture medium by pipetting. Culture the isolated clones for 2–3 days until most clones become confluent.
6. Duplicate the 96-well plate after trypsinization, and culture the cells on the plates for an additional 2–3 days.

One 96-well plate will be used to prepare a frozen stock and the other will be used to prepare genomic DNA for PCR genotyping.

### 4.2. Frozen mini-stock

1. Remove the medium from one 96-well plate and wash cells once with 100 μL of PBS. Add 10 μL of trypsin/EDTA and incubate at 37 °C for 3 min.
2. Resuspend cells in 50 μL of medium and transfer them to 0.75 μL Matrix Storage Tubes (ThermoFisher Scientific) containing 50 μL of Bambanker DIRECT medium (Nippon Genetics, CS-06-001). Mix the cells by pipetting.
3. Close the tubes with a cap and store them at −80 °C until PCR genotyping is finished.

### 4.3. Genomic DNA preparation

1. Remove the medium from the 96-well plate and wash cells once with 100 μL of PBS. Add 75 μL of the DirectPCR working solution (0.5 × DirectPCR Lysis Reagent-Cell [Viagen Biotech, 302-C] containing 0.5 mg/ml of Proteinase K).
2. Seal the plate with an aluminium seal and incubate it at 55 °C for >6 h in a rocking incubator.
3. Place the plate in a Tupperware containing wet tissue papers and float the wet chamber in a water bath at 85 °C for 1.5 h, to inactivate proteinase K.
4. Spin the plate to collect the samples and use 1 μL for PCR genotyping.

### 4.4. Genotyping by PCR

1. Design appropriate primers to check the insertion by PCR (**Figure 4A** and **B**). We designed a primer set to detect both the wild-type (WT) and inserted alleles (primer set a), and another primer set to detect only the inserted allele (primer set b).
2. Set up a PCR reaction using Tks Gflex DNA Polymerase (Takara Bio, R060A) (1 × Gflex PCR Buffer, 0.3 μM primers and 0.4 U of Tks Gflex DNA Polymerase in a 15 μL reaction). Add 1 μL of the genomic DNA to the 15 μL reaction mixture. As controls, add genomic DNA extracted from the parental cells to the reaction mixture. Perform PCR (30 cycles of 98 °C for 10 s, 55 °C for 15 s and 68 °C for 0.5 min/kb).
3. Examine PCR products using agarose gel electrophoresis or the MultiNA microchip electrophoresis system (Shimadzu MCE-202) (**Figure 4C**).
4. Identify clones showing the expected band patterns of biallelic insertion. An example of N-terminal tagging of CENPC is shown in **Figure 4C**. Clones with biallelic insertion are highlighted by a red circle (clones 2 and 8). These clones can be checked further by genomic sequencing. Expand these clones of the mini-stock kept at −80 °C.

**Figure 4.**
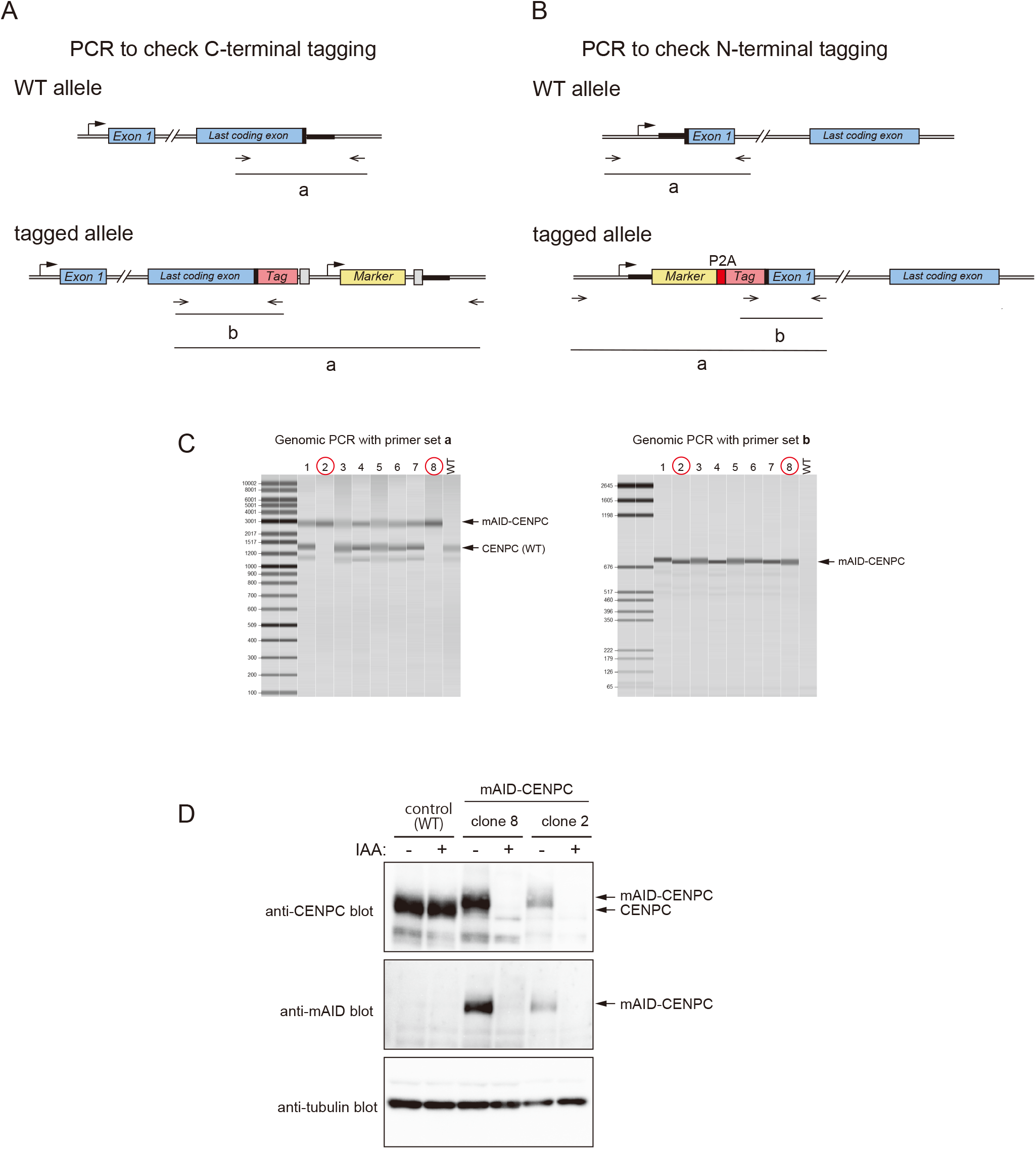
Confirmation of isolated clones. (**A**) Genotyping of C-terminally tagged alleles by PCR. The primer set (a) amplifies a smaller PCR product (1–1.5 kb) from the WT allele, while it amplifies a larger product (1–1.5 kb plus the size of the insertion) from the tagged allele. The primer set (b) generates a PCR product from the tagged allele exclusively (and not from the WT allele). Note that the primer set (a) must be designed outside of the homology arms. (**B**) Genotyping of N-terminally tagged alleles by PCR. The PCR strategy is analogous to that used to analyse C-terminal tagging. (**C**) PCR genotyping of the CENPC allele, in which a Hygro-P2A-mAID-mClover cassette was inserted at the N-terminal coding region. HCT116 CMV-OsTIR1 parental cells were used. (**D**) Confirmation of the fusion protein by WB. WT or mAID-CENPC clones in the HCT116 CMV-OsTIR1 background were treated with DMSO or 500 μM IAA for 24 h. Anti-CENPC, anti-mAID and anti-tubulin antibodies (MBL, PD030, M214-3 and M175-3, respectively) were used for detection.

## 5. Testing degradation of mAID-fused proteins

If mAID-mClover or mAID-mCherry2 was fused to a POI, depletion of the mAID-fused POI upon the addition of auxin can be checked easily by flow cytometry or microscopy. If an antibody that detects the POI is available, it is possible to assess the loss of the WT protein and confirm the expression of the mAID-fused protein. Next, we describe our methods to check clones by Western blotting and flow cytometry.

### 5.1. Confirmation by Western blotting

1. Grow clones that have been confirmed by PCR in a 6-well plate. To prepare a control with mock treatment (DMSO), one clone should be grown in two wells. Grow cells to 50% confluency.
2. Prepare a 500 mM stock solution of IAA in DMSO and keep aliquots at −20 °C.
3. If testing cells constitutively expressing OsTIR1 (CMV-OsTIR1), add IAA to one well at the final concentration of 200–500 μM. Cells growing in the other well should be mock treated with DMSO. To test cells conditionally expressing OsTIR1 (Tet-OsTIR1), add IAA and 0.2 μg/mL of doxycycline. Grow the treated cells for 12–24 h.
4. Collect cells by trypsinization and wash them in culture media.
5. Collect cells in a microtube and wash once with PBS.
6. Mix the cell pellet in 50 μL of RIPA buffer (25 mM Tris-HCl [pH 7.6], 150 mM NaCl, 1% NP40, 1% sodium deoxycholate and 0.1 % SDS) and incubate the tube for 30 min on ice.
7. Remove debris by centrifugation at 10000 rpm for 5 min at 4 °C. Collect the supernatant and mix with the same volume of 2 × Laemmli SDS sample buffer (Tris-HCl [pH 6.8], 4% SDS, 20% glycerol, 10% 2-mercaptoethanol and 0.004% bromophenol blue) before incubating at 95 °C for 5 min.
8. Typically, load 5 μL of the protein sample for Western blotting. Commercial antibodies are available to detect mAID and OsTIR1 (MBL M214-3 and PD048, respectively). The example of mAID-fused CENPC is shown in **Figure 4D**.

### 5.2. Confirmation by flow cytometry

1. Grow and treat cells in a 6-well plate as described in section 5.1.
2. Remove the media and wash cells once with PBS. After trypsinization, resuspend cells in media and transfer them to a microtube. Remove most of the media after spinning down cells and resuspend them in the residual medium by pipetting.
3. Resuspend the cells in 1 mL of 4% paraformaldehyde/PBS by pipetting. Incubate in the dark at 4 °C. Note that a methanol-free paraformaldehyde solution should be used to preserve the fluorescence of mClover and mCherry2.
4. Before flow cytometric analysis, wash the fixed cells once with PBS containing 1% of BSA. Resuspend the washed cells in an appropriate volume of PBS containing 1% of BSA and analyse them after filtration with a strainer to remove aggregation (An example of flow cytometry data is shown in **Figure 5** and **Figure 6**).

**Figure 5.**
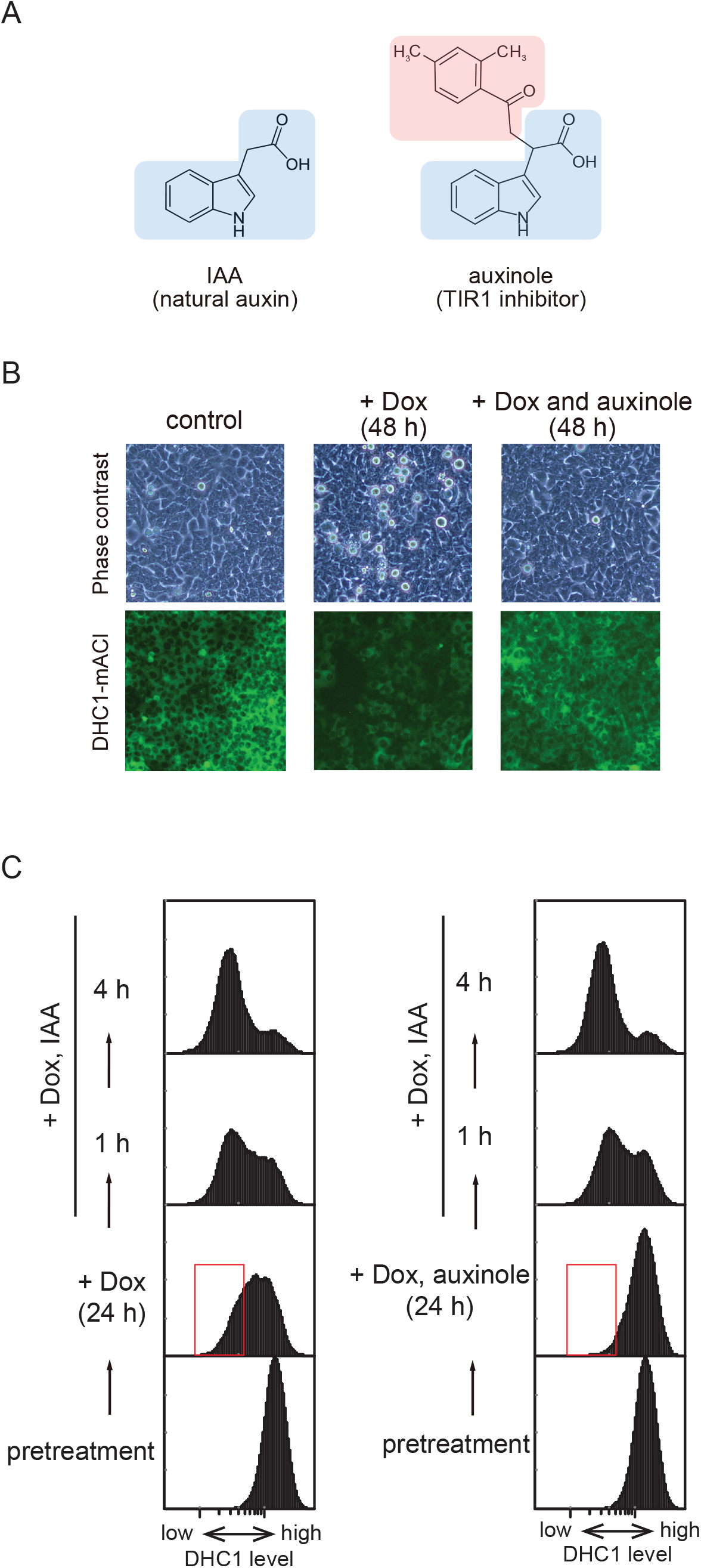
Use of auxinole to control DHC1-mAID-mClover (DHC1-mACl) Tet-OsTIR1 cells. (**A**) The structure of IAA and auxinole. (**B**) Microscopic analysis of DHC1-mAC Tet-OsTIR1 cells. The cells were treated with 0.2 μg/mL of doxycycline (Dox) or Dox with 200 μM auxinole for 48 h before microscopy. (**C**) Flow cytometric analysis of DHC1-mAC Tet-OsTIR1 cells. The cells were treated with 0.2 μg/mL of Dox or Dox with 200 μM auxinole for 24 h before replacing the culture medium with Dox and 500 μM IAA.

**Figure 6.**
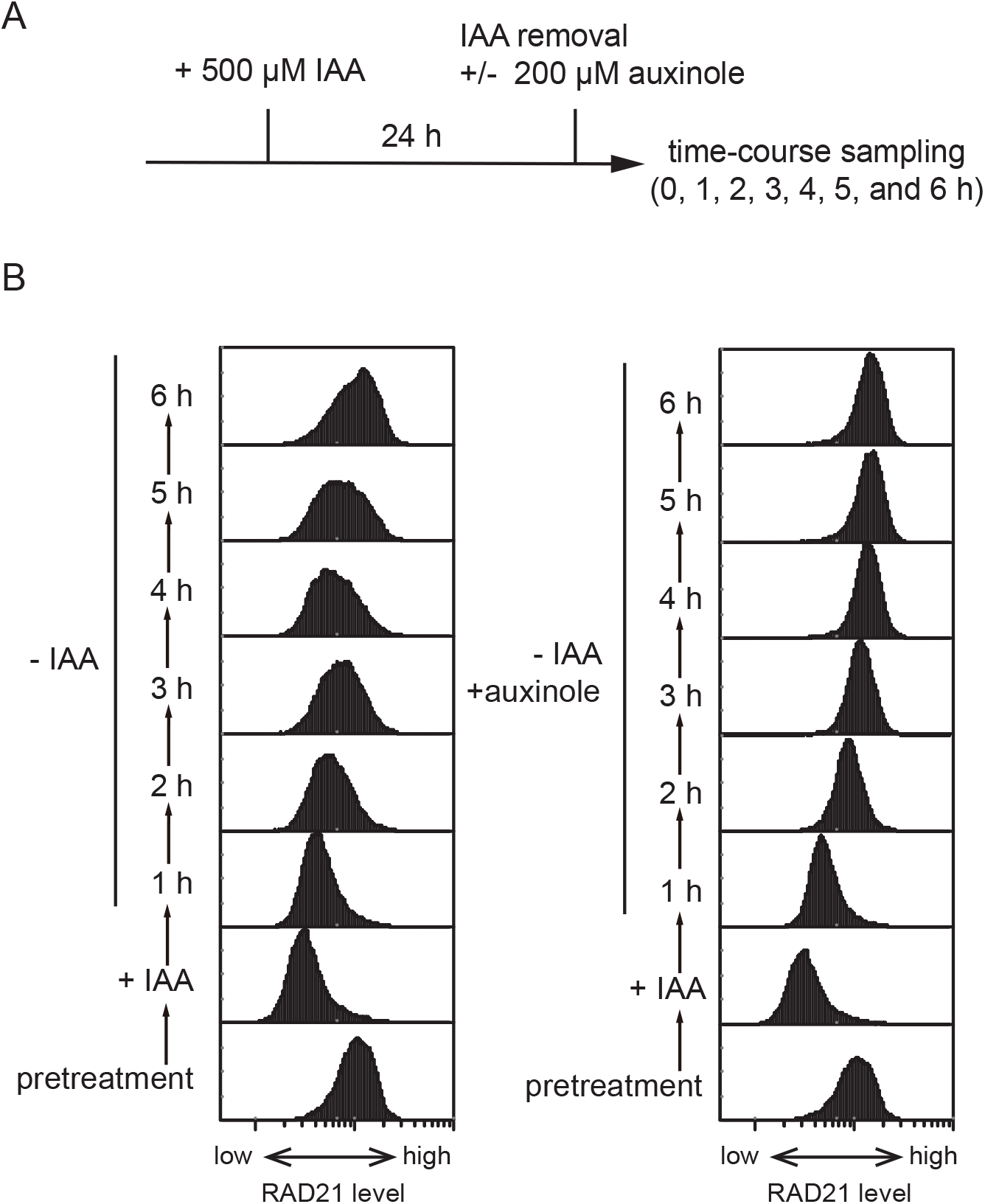
Re-expression of RAD21-mAID-mClover (RAD21-mACl). (**A**) Scheme of the experiment. The RAD21-mACl CMV-OsTIR1 cells were treated with 500 μM IAA for 24 h before replacing the medium with fresh medium with or without 200 μM auxinole. (**B**) Flow cytometric analysis of RAD21-mACl cells.

## 6. Suppression of degradation using auxinole

We demonstrated previously that HCT116 cells in which mAID-mClover was fused to the cytoplasmic dynein heavy chain 1 (DHC1) protein showed a growth defect when OsTIR1 was expressed without IAA [6]. This was caused by partial degradation of DHC1-mAID-mClover (DHC1-mACl). We subsequently noted that, in some cases, the expression level of mAID-fused proteins was significantly reduced in cells constitutively expressing OsTIR1 (data not shown). Moreover, in some cases, we could only introduce a biallelic mAID into a POI in the conditional Tet-OsTIR1 background (and not in the constitutive CMV-OsTIR1 background) [6]. These results indicated that mAID-tagged proteins could be ubiquitylated by OsTIR1 for partial degradation even without the addition of exogenous IAA.

We aimed to suppress this basal degradation by using a chemical antagonist of TIR1. Previously, we developed a potential TIR1 inhibitor named auxinole to control auxin signalling in plants [16]. Auxinole contains a moiety that is analogous to IAA and binds to the IAA-binding pocket of TIR1 (**Figure 5A**, shown in blue). Because of steric inhibition by the additional dimethylphenylethyl-2-oxo moiety (**Figure 5A**, shown in red), auxinole blocks the association of the degron domain of the AUX/IAA proteins. For the experiments in this work, auxinole was dissolved in DMSO to make a 200 mM stock solution, which was stored at −20 C° (commercially available from BioAcademia #30-001).

We initially tested whether auxinole affected the growth of HCT116 cells and found that it did not alter the cell cycle or colony formation when added up to 200 μM (data not shown). Subsequently, we tested the suppression of basal degradation using DHC1-mACl cells in the HCT116 Tet-OsTIR1 background [6]. Addition of doxycycline induced OsTIR1 expression, which caused partial depletion of DHC1-mACl and mitotic arrest in many cells, as we reported previously (**Figure 5B**) [6]. This showed that OsTIR1 expression, even in the absence of auxin, induced a mitotic phenotype that was analogous to knockdown or inhibition of dynein [19, 20]. The addition of auxinole together with doxycycline clearly suppressed the downregulation of DHC1-mACl and the mitotic arrest (**Figure 5B**). To test whether DHC1-mACl could be rapidly depleted, we added doxycycline with or without auxinole for 24 h. We monitored the expression levels of DHC1-mACl by flow cytometry, and found that basal degradation was mostly suppressed in the cells treated with doxycycline and auxinole (**Figure 5C**, compare the boxes shown in red). Subsequently, the culture media was replaced with fresh one containing doxycycline and IAA, but not auxinole. **Figure 5C** shows that DHC1-mACl was rapidly degraded after medium replacement and was mostly depleted within 4 h.

An advantage of AID technology is that the expression level of mAID-fused proteins can be reversibly controlled [2]. We expected that auxinole would be useful for re-expression after depletion, because IAA-bound OsTIR1 can remain active for a while, even after the removal of IAA from the culture medium. To test this idea, we used HCT116 CMV-OsTIR1 cells in which the cohesin subunit RAD21 was fused to mAID-mClover (RAD21-mACl) [6]. Initially, we depleted RAD21-mACl by adding IAA for 24 h (**Figure 6A**). Subsequently, we replaced the medium with fresh media with or without auxinole, and collected time-course samples to monitor the expression levels of RAD21-mACl by flow cytometry (**Figure 6B**). We found that recovery of RAD21-mADl was significantly rapid and sharp when auxinole was added, compared with cells without auxinole. These results suggest that the OsTIR1 inhibitor auxinole is useful for the tight control of the expression of mAID-fused proteins in human cells.

## 7. Conclusion

We described a CRISPR–Cas9-based method that can be used to fuse endogenous POIs to mAID and other tags. We developed new plasmids for N-terminal tagging, so that it is now possible to tag the C and N termini of POIs (**Figure 2**). To suppress basal degradation in cells expressing OsTIR1, we used the OsTIR1 antagonist auxinole (**Figure 5A**). Even in the Tet-OsTIR1 background cells, it is now possible to induce rapid degradation of mAID-fused proteins by inducing OsTIR1 in the presence of auxinole (**Figure 5C**). Moreover, auxinole is useful for the re-expression of mAID-fused POIs after depletion (**Figure 6B**). The use of auxinole allows the rapid, tight and efficient control of the expression of mAID-fused POIs. Other genetic systems also enable the control of degron- or tag-fused POIs using a chemical ligand [1, 21–23]. However, to the best of our knowledge, there are no inhibitors that allow the tight control of these systems. AID technology—now combined with the degradation inducer, auxin, and the inhibitor, auxinole—will be particularly useful to dissect biological networks, such as transcriptional cascades, signal transduction, and cell-cycle control systems, in which a primary defect caused by the loss of a POI leads to secondary defects. We hope that the method described in this manuscript will enhance the utility of AID technology for functional studies of endogenous proteins in living cells.

## Acknowledgements

We thank Venny Santosa, Akemi Mizuguchi and Tomoko Ashikawa for their support. This study was supported by JSPS KAKENHI grants 17K15068 to TN, and 18H02170 and 18H04719 to MTK; and by research grants from JST A-STEP (grant number AS2915150U), the Canon Foundation, the Asahi Glass Foundation, and the Takeda Science Foundation.

