## Supplementary Figures for "Generation of conditional auxin-inducible degron (AID) cells and tight control of degron-fused proteins using the degradation inhibitor auxinole"

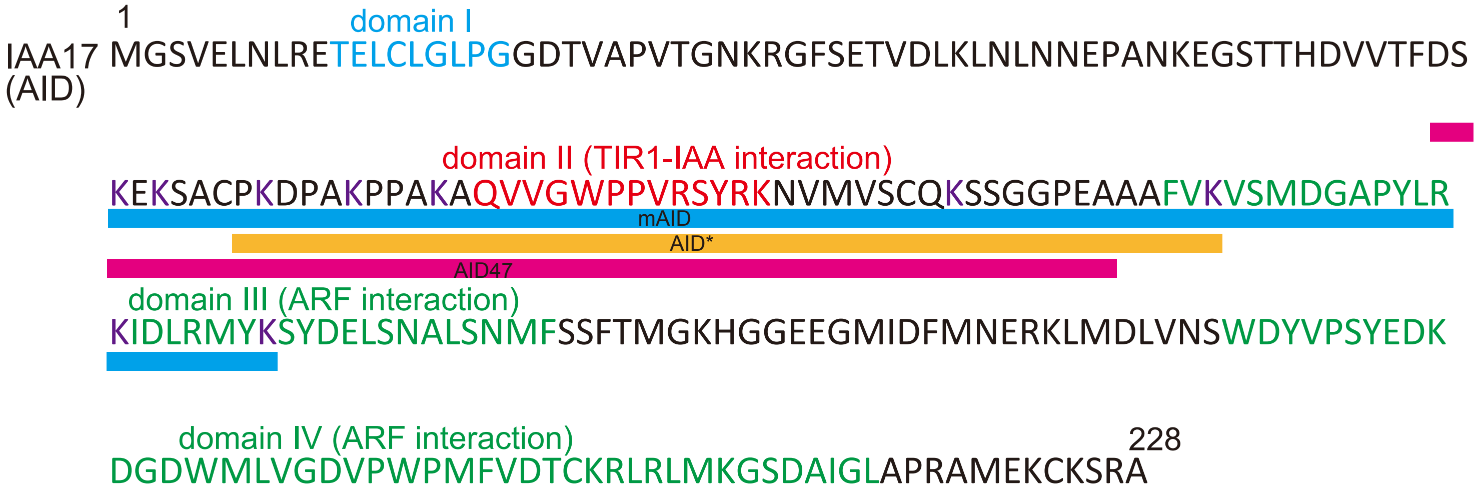


Supplementary Figure 1
The amino-acid sequence of IAA17 (AID) and truncated AID variants (mAID, AID*, and AID47). The domains I to IV are highly conserved among the AUX/IAA-family proteins. The domain II (shown in red) is the essential region required for the interaction with TIR1-IAA. The domains III and IV (shown in green) are responsible for the interaction with Auxin-Response Factors (ARFs).  Lysine resides within mAID are colored in purple.

Plasmids for making parental cells expressing OsTIR1

| Plasmid name | Transgene | Marker |
|---|---|---|
| pMK232 | CMV-OsTIR1 | Puro |
| pMK243 | Tet-OsTIR1 | Puro |
| pMK364 | CMV-OsTIR1 | loxP-Puro-loxP |
| pMK365 | Tet-OsTIR1 | loxP-Puro-loxP |

| Plasmid name | Transgene |
|---|---|
| AAVS1 T2 CRISPR in pX330 | spCas9, AAVS1 T2 gRNA |

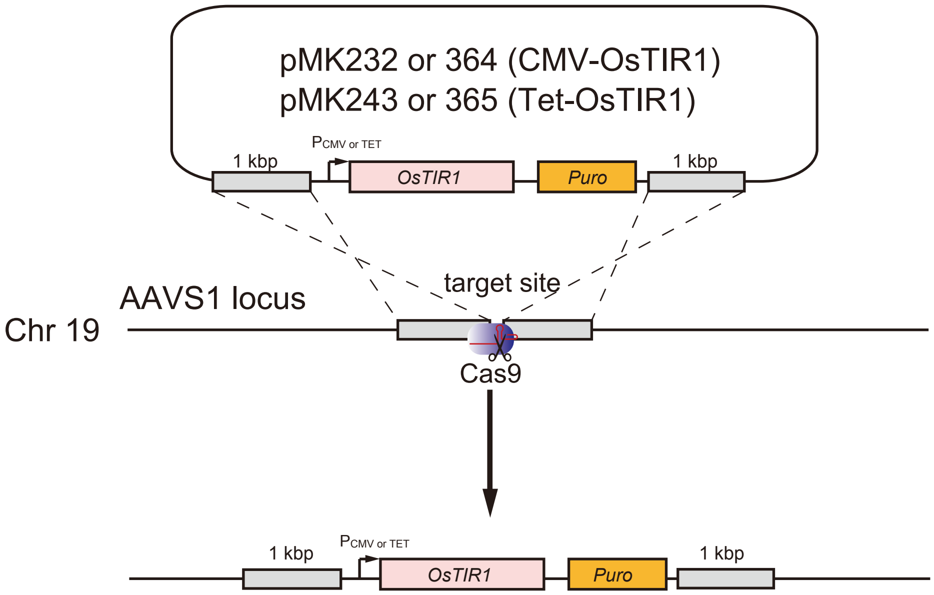


Supplementary Figure 2

List of plasmids for introducing OsTIR1 and illustration of the integration at the AAVS1 locus .
